# γH2AX in the S Phase After UV Irradiation Corresponds to DNA Replication and Not to the Extent of DNA Damage

**DOI:** 10.1101/810689

**Authors:** Shivnarayan Dhuppar, Sitara Roy, Aprotim Mazumder

## Abstract

Ultraviolet (UV) radiation is a major environmental mutagen. Exposure to UV leads to a sharp peak of γH2AX – the phosphorylated form of a histone variant H2AX – in the S phase within an asynchronous population of cells. γH2AX is often considered as a definitive marker of DNA damage inside a cell. In this report we show that γH2AX in the S phase cells after UV irradiation does not report on the extent of primary DNA damage in the form of cyclobutane pyrimidine dimers or on the extent of its secondary manifestations as DNA double strand breaks or in the inhibition of global transcription. Instead γH2AX in the S phase corresponds to the sites of active replication at the time of UV irradiation – despite which, the cells complete the replication of their genomes and arrest within the G2 phase. Moreover, cells in all the phases of the cell cycle develop similar levels of DNA damage. Our study suggests that it is not DNA damage but the response elicited, which peaks in the S phase upon UV damage.

## Introduction

Chromatin, a complex of DNA and protein, transforms DNA inside a nucleus into a functional genome. It has histones which make up the majority of its protein content and whose modifications in the form of histone methylation, -acetylation or -phosphorylation are major regulators of nuclear processes such as DNA transcription, -replication or -repair [1, 2]. One such histone modification is called γH2AX: the phosphorylation of a variant histone H2AX at serine 139 observed during a variety of cellular stresses [3–5]. Genotoxic stress, in particular, leads to the accumulation of γH2AX at the sites of DNA damage within just a few minutes of damage which at later timepoints starts to spread across mega base-pairs from those sites of damage [6]. This phosphorylation of H2AX at serine 139 is mediated directly by the phosphatidylinositol 3-kinase-related kinases (PIKK) ataxia telangiectasia mutated (ATM) and ataxia telangiectasia and Rad3-related (ATR)—the master transducers of DNA signals [7]—which has established γH2AX as a standard, direct and faithful marker of DNA damage inside a cell [6]. Using the levels of γH2AX as a measure of DNA damage, in particular that of DNA double strand breaks (DSBs), previous studies have shown that DNA damage peaks in the S phase of the cell cycle with a variety of genotoxic treatments [8–11]. These observations conform with the idea that S phase cells within a population are most vulnerable to DNA damage owing to the active replication of their DNA which requires permissible chromatin states that render the DNA more susceptible to DNA damage [12], or the fact that single strand breaks (SSBs) formed as repair intermediates may be converted to DSBs upon unchecked replication [13]. This notion though has been challenged by a relatively recent study where it has been shown that only a minority of γH2AX foci in the S phase after UV irradiation represent DSBs [14]:—the study while addressing many important aspects of UV-induced DNA damage, does not comment on the nature of γH2AX in the S phase cells apart from the observation that it does not mark DSBs.

Here we fill the above lacuna by studying cell cycle-dependent DNA damage responses (DDR) using an improved imaging-based cell cycle staging as developed previously [8]. This helped us study not only the levels of different DDR proteins at a single-cell resolution but also their localization across the cell cycle. In particular, we show that the γH2AX peak in the S phase after UV irradiation reflects the colocalization of γH2AX with the active replication forks at the time of irradiation and reports neither on the extent of primary DNA damage as measured by the induction of UV adducts, nor on that of the secondary damage reflected by the DSBs in those cells. Moreover, we show that this colocalization of γH2AX with the sites of active replication does not entirely halt replication and that most of the S phase cells at the time of UV irradiation complete the replication of their DNA and are later arrested in the G2 phase of the cell cycle. This further proves that cells in all the cell cycle phases are equally vulnerable to DNA damage from external sources, but it is the S phase cells which show the highest levels of response in terms of γH2AX induction owing to active DNA replication in those cells.

## Results

### Cell cycle-dependent DNA damage responses

Cell cycle is the process by which a cell divides into two daughter cells. Starting from the first gap phase G1 the cell moves onto the S phase where it replicates its DNA and ends up in the second gap phase G2 which then is followed by the cell division in the mitosis or M phase. Each cell cycle phase is marked by a distinct chromatin state facilitating the structure, function and expression of the genome in that particular phase of the cell cycle [2, 15, 16]. S phase, in particular, requires an open, more permissive chromatin state to aid DNA replication. This opened-up chromatin state, it is believed, renders the S phase cells more vulnerable to DNA damage [12]. Here we investigated the above hypothesis by studying cell cycle-dependent DNA damage responses (DDR) to UV and neocarzinostatin (NCS) treatments. UV is known to cause replication stress by forming cyclobutane-pyrimidine dimers and 6-4 pyrimidine-pyrimidone photoproducts that are repaired by Nucleotide Excision Repair (NER), while NCS—a radiomimetic drug—causes DSBs within just a few minutes of the treatment [17] that may be repaired by Homologous Recombination (HR) or Non-homologous End Joining (NHEJ) [18, 19]. The aim was to study differences and similarities between the responses to the two treatments, especially in the S phase of the cell cycle.

DNA content-based cell cycle staging may lead to mislabelling of early S phase cells as G1 or late S phase cells as G2 depending on their degree of progress inside the S phase. Hence to identify true S phase cells in the population we labelled them with a thymidine analogue 5-ethynyl-2’-deoxyuridine (EdU). EdU gets incorporated into the actively replicating DNA, which can later be detected via click chemistry [20]. By combining EdU-based labelling of the S phase cells with a previously-developed microscopy-based cell cycle staging from DNA content [8] we were able to identify true S phase cells which previously were mislabelled as G1 or G2 phase cells based just on their DNA contents (Figures 1A and 1B). Once standardized we used the technique to first study the DDR in the S phase cells when treated with UV and NCS in terms of the levels of phosphorylation of the important DDR proteins ATM and H2AX.

**Figure 1:**
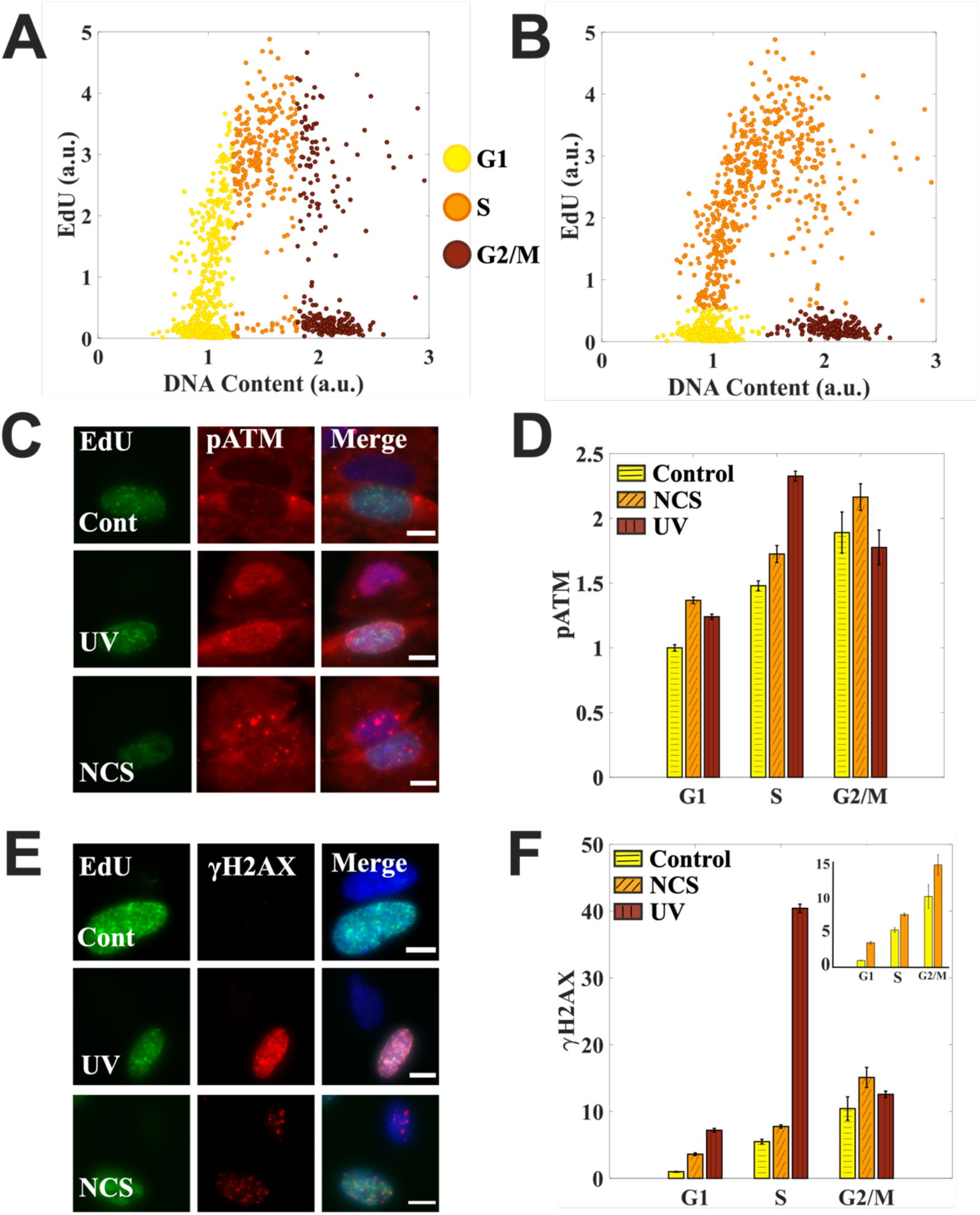
Cell cycle-dependent DNA damage responses. (A) Cell cycle staging based on just the DNA content. (B) The same cells as in (A) are staged also according to their EdU content. EdU helps identify true S phase cells. (C) Cell cycle-dependent induction of pATM after UV and NCS treatment. Each row has two cells: one S phase (EdU positive), other non-S phase. (D) Quantification for (C). UV-treated cells have pATM peak in the S phase while NCS treated cells have pATM levels increase with the increase in DNA content as moving from G1 to S to G2/M. (E) Cell cycle-dependent induction of γH2AX after UV and NCS treatment. Each row has two cells: one S phase, other non-S phase. (F) Quantification for (E). UV-treated cells have a sharp γH2AX peak in the S phase while NCS treated cells have pATM levels increase with the increase in DNA content as moving from G1 to S to G2/M. Inset compares just the control and NCS-treated cells. Cells were treated with 10 J/m^2^ and 1.6 μg/ml NCS for two minutes. Bar graphs are normalized with respect to the mean value for G1 phase in control cells across the populations. Scale bar: 10 μm

ATM is an important mediator of DNA damage response in its active form which is marked by its phosphorylation at serine 1981 (pATM). It is activated during both NCS (which causes DSBs) and UV (which causes replication stress) treatments but via different mechanisms: DSBs directly recruit ATM via MRN complex which leads to autophosphorylation of ATM while UV causes ATM activation in an ATR-dependent manner [21]. Once activated ATM also can phosphorylate H2AX at serine 139—a modification known by the name γH2AX and considered as a standard DNA damage marker.

We observed that for the cells treated with NCS the mean levels of pATM and γH2AX increased with the increase in DNA content as moving from G1 to S to G2/M phases of the cell cycle. While for the cells treated with UV the mean levels of both pATM and γH2AX peaked in the S phase of the cell cycle with γH2AX peak being much more pronounced than the pATM peak (Figures 1D and 1F). The above observations suggest two possibilities: 1) There is no distinction among the cell cycle phases as to the extent of primary DNA damage. Rather it increases with the increase in DNA content as seen in NCS treated cells, or 2) The S phase cells are more vulnerable to DNA damage as inferred from the large γH2AX peak after UV irradiation.

While NCS treatment leads to a clear accumulation of γH2AX at the sites of DSBs within a few minutes of the treatment, the role of γH2AX in the S phase cells post UV irradiation is not well known. Hence to evaluate the above two possibilities we started out by first investigating the cell cycle profile of UV-induced primary DNA lesions.

### UV-induced primary DNA lesions do not peak in the S phase unlike the DNA damage response elicited in terms of γH2AX

UV irradiation induces direct photo-damage to DNA in the form of cyclobutane pyrimidine dimers (CPDs) and 6-4 pyrimidine-pyrimidone photoproducts of which CPDs constitute up to 80% of the total photoproducts [22]. We stained cells for CPDs along with γH2AX to measure the extent of primary DNA damage along with that of the response elicited on a cell-by-cell basis in the context of cell cycle (Figure 2A).

**Figure 2:**
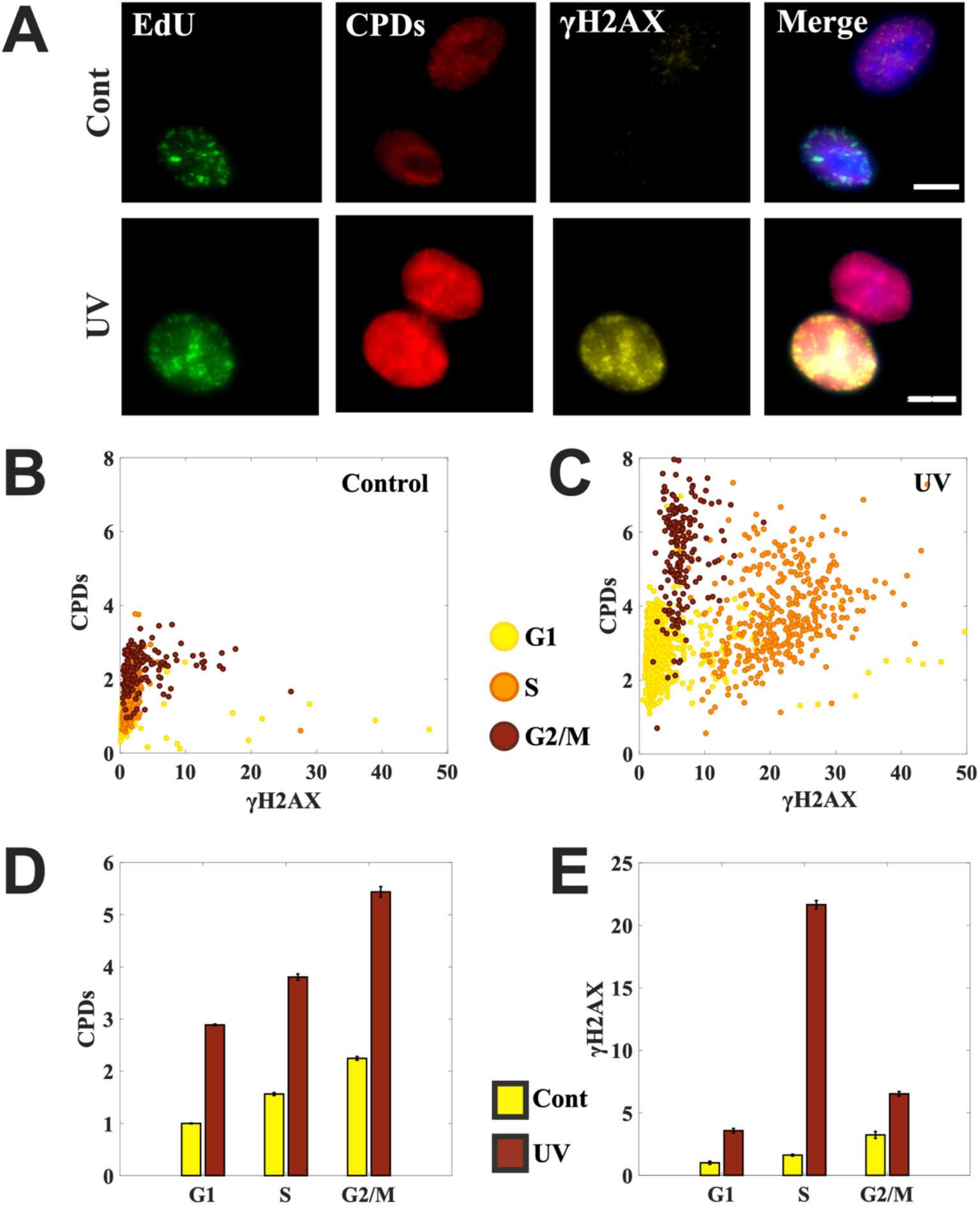
UV-induced primary DNA lesions do not peak in the S phase unlike the DNA damage response elicited in terms of γH2AX. (A) Control and UV-treated cells were stained simultaneously for cyclobutane pyrimidine dimers (CPDs) and γH2AX to measure the extent of primary DNA damage (CPDs) and compare response (γH2AX) in S and non-S phase cells. Each row has two cells: one S phase (EdU positive), other non-S phase. (B, C) Scatter plots for control and UV-treated cells marking the levels of primary damage in the form of CPDs and the response it elicited in the form of γH2AX induction at single-cell resolution. UV irradiation gives rise to 3 distinct populations corresponding to G1, S and G2/M phases in terms of the levels of primary DNA damage and the response it elicits. (D, E) Bar graphs comparing the mean levels of primary damage in the form of CPDs in control and UV-treated cells and the response in the form of γH2AX. Mean levels of CPDs increase with the increase in DNA content as moving from G1 to S to G2 phases of the cell cycle. Moreover, the S phase peak observed for γH2AX is missing for the levels of primary damage in the form of CPDs. Cells were irradiated with 10 J/m^2^. Scale bar: 10 μm.

We observed that UV irradiation gives rise to three distinct population of cells corresponding to G1, S and G2/M phases on the basis of the levels of CPDs and that of γH2AX as reflected in the single cell scatter plots (Figure 2B and 2C). Moreover, γH2AX is markedly higher in the S phase cells of the UV-treated population, while such distinction is not observed in terms of CPD levels (Figure 2C). In fact, mean levels of CPDs in UV treated population increase with the increase in DNA content as moving from G1 to S to G2 phases of the cell cycle while mean γH2AX levels show a clear peak in the S phase of the cell cycle (Figure 2D and 2E). Furthermore, γH2AX foci do not correspond to regions of high CPD intensity (Figure 2A).

The above observations on UV-induced DNA damage clearly show that it is not DNA damage itself, but the response it elicits, which is cell cycle-dependent with a clear peak in the S phase. Moreover, these observations also suggest that UV-induced γH2AX in the S phase does not correspond to primary DNA lesions from UV. Thus it is not that the permissive chromatin state of the S phase cells result in increased UV damage, which causes a standard marker of DDR like γH2AX to peak in this phase. But instead of the primary UV adducts, could it be the DSBs formed from secondary repair intermediates like SSBs during replication, that cause the γH2AX peak? To address this we investigate the colocalization of the γH2AX foci with a standard marker of DSBs in the next section.

### UV-induced γH2AX does not report also on the extent of secondary DNA damage in the S phase cells

UV-induced DNA adducts can cause replication fork stalling and are repaired by nucleotide excision repair that results in single strand breaks (SSBs) as repair intermediates. These repair intermediates, if unrepaired, can lead to deleterious DSBs upon DNA replication [13]. 53BP1 is a key component of double strand break repair pathways and like γH2AX is known to accumulate at the sites of DSBs within a few minutes of damage [23, 24]. It forms distinct foci at the sites of DSBs where it colocalizes with γH2AX and has been used previously to count the number of DSBs in the cells [17, 25]. We used a similar metric whereby if there is more than 50% colocalization between γH2AX and 53BP1 foci at a site, then the site is considered as a DSB site (Figure 3A). We used this metric to calculate the percentage of γH2AX foci that corresponds to DSBs in the S phase cells post UV.

**Figure 3:**
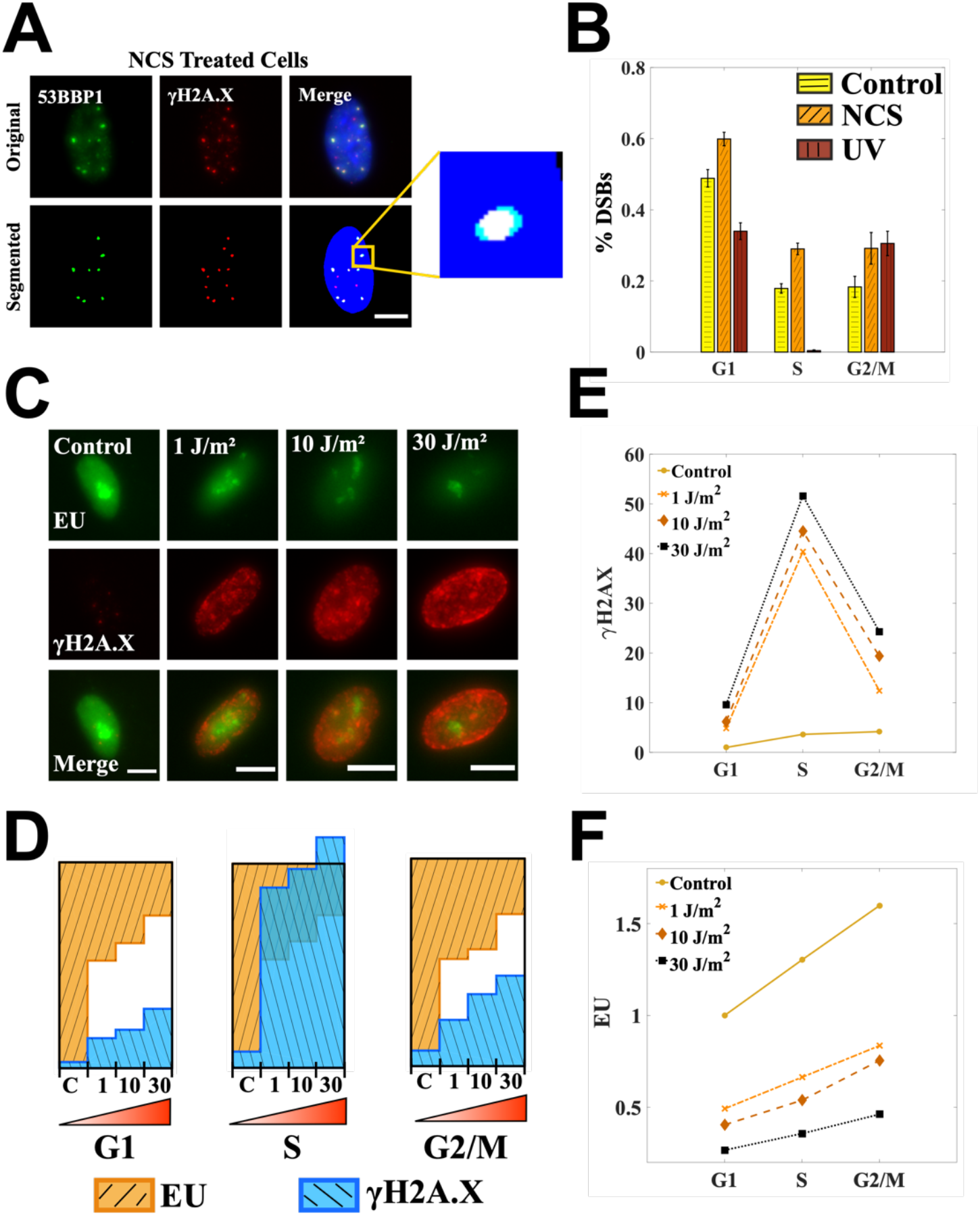
UV-induced γH2AX does not report also on the extent of secondary DNA damage in the S phase cells. (A) Definition of the colocalization metric: more than 50% overlap between the foci from the two channels at a position. Images are of NCS-treated cells: those 53BP1 foci colocalizing with γH2AX foci are considered as DSBs (B) Percentage of γH2AX foci which are double strand breaks for control, UV- and NCS-treated cells. For UV treatment a very little fraction of γH2AX foci in the S phase cells corresponds to DSBs. (C) Global transcription is measured in UV-treated cells by quantifying 5-ethynyl-uridine incorporation. (D) Bar graphs for γH2AX and EU are normalized across the four dosages. Resultant bar graphs for EU are inverted over those for γH2AX such that the total height of the two bars corresponding to control population in every cell cycle phase is unity. A clear gap is observed between γH2AX and EU bars for G1 and G2/M phases after UV treatment. The γH2AX bars in the S phase not just overlap with EU bars but goes past the unit box showing a disproportionate increase in γH2AX. (E) Mean levels of γH2AX increase with the increase in DNA damage as achieved by increasing the UV dosage in all the phases of the cell cycle. For all the dosages, there is a sharp γH2AX peak in the S phase. (F) Increase in DNA damage always leads to decrease in EU as seen across the population treated with different dosages of UV. But within a population there is no dip in EU levels corresponding to the γH2AX peak showing that γH2AX in the S phase after UV does not reflect on the total extent of DNA damage in those cells. Cells were treated with 10 J/m^2^ and 1.6 μg/ml NCS for two minutes. Cells were let to recover for 30 minutes in the presence of EU. Line graphs are showing relative levels with respect to the mean value of G1 phase in control cells. Scale bar: 10 μm.

We found that for the cells treated with UV the percentage of γH2AX foci representing DSBs was remarkably small in the S phase cells unlike the large percentage observed in the cells treated with a direct double strand breaks-causing agent NCS (Figure 3B). The observation conforms with a previous study showing that only a minority of γH2AX foci after UV irradiation contain double strand breaks [14]. With DSBs out of consideration, we wondered what other kinds of DNA damage could the UV-induced γH2AX in the S phase represent. Instead of going one-by-one through all possible kinds of damage that UV can possibly cause, we used a proxy that can report on total levels of DNA damage inside the cells fairly accurately—the comparative levels of total transcriptional activity inside a cell [26, 27].

UV-induced DNA damage repair depends on the transcriptional status of the damaged DNA: an actively transcribed strand could be repaired via transcription-coupled NER while the rest of the DNA could repaired via global genomic NER [28]. This implies that transcription is a direct target of damage repaired by NER processes and that the total transcriptional activity inside a cell can potentially report on the extent of DNA damage inside a cell. Moreover, inhibition of global transcription is one of the first few responses to UV irradiation [29, 30]. In fact, there have been studies reporting inhibition of global transcription as a proxy for DNA damage, especially from UV exposure [26, 27]. We measured the levels of global transcription by labelling the nascent RNA with 5-ethynyl-uridine (EU) which can later be detected via click chemistry [31].

We observed that an increase in DNA damage by increasing the UV dosage always led to a decrease in global transcription irrespective of the cell cycle phase (Figure 3F). In fact the fold decrease in global transcription across the population was almost the same in all the three phases of the cell cycle (Bar graphs for EU in Figure 3D). But unlike G1 and G2/M phases of the cell cycle, the increase in γH2AX levels in the S phase cells was disproportionately high and was not reflected in the extent of transcriptional inhibition in that phase across the population (Figure 3D). Despite having the highest levels of γH2AX within a population, the S phase cells did not show a corresponding dip in the global transcription as one would expect if γH2AX levels in the S phase were indeed reporting on the extent of DNA damage in those cells (Figures 3E and 3F).

Taken together the above observations suggest that all of γH2AX in the S phase cells after UV irradiation cannot be due to DNA damage. To further investigate the nature of γH2AX in the S phase cells after UV irradiation we looked at its localization with respect to replication forks in those cells as discussed next.

### γH2AX colocalizes with the sites of active replication on UV irradiation

The spatial resolution inbuilt within the imaging-based cell cycle staging helped us look at the localization of γH2AX in the nucleus after genotoxic treatments. We observed that γH2AX on UV irradiation colocalized with the sites of active replication unlike for NCS treatment where it colocalized specifically to the sites of DSBs. The percentage of γH2AX foci colocalizing with the sites of active replication as marked by EdU incorporation was more than 60% for UV-irradiated cells while it was smaller than 3% for control and NCS-treated cells (Figures 4A and 4B). This accumulation of γH2AX at the sites of replication started to build up within a few minutes of exposure and peaked at around 30 minutes post UV irradiation (Figure 4C).

**Figure 4:**
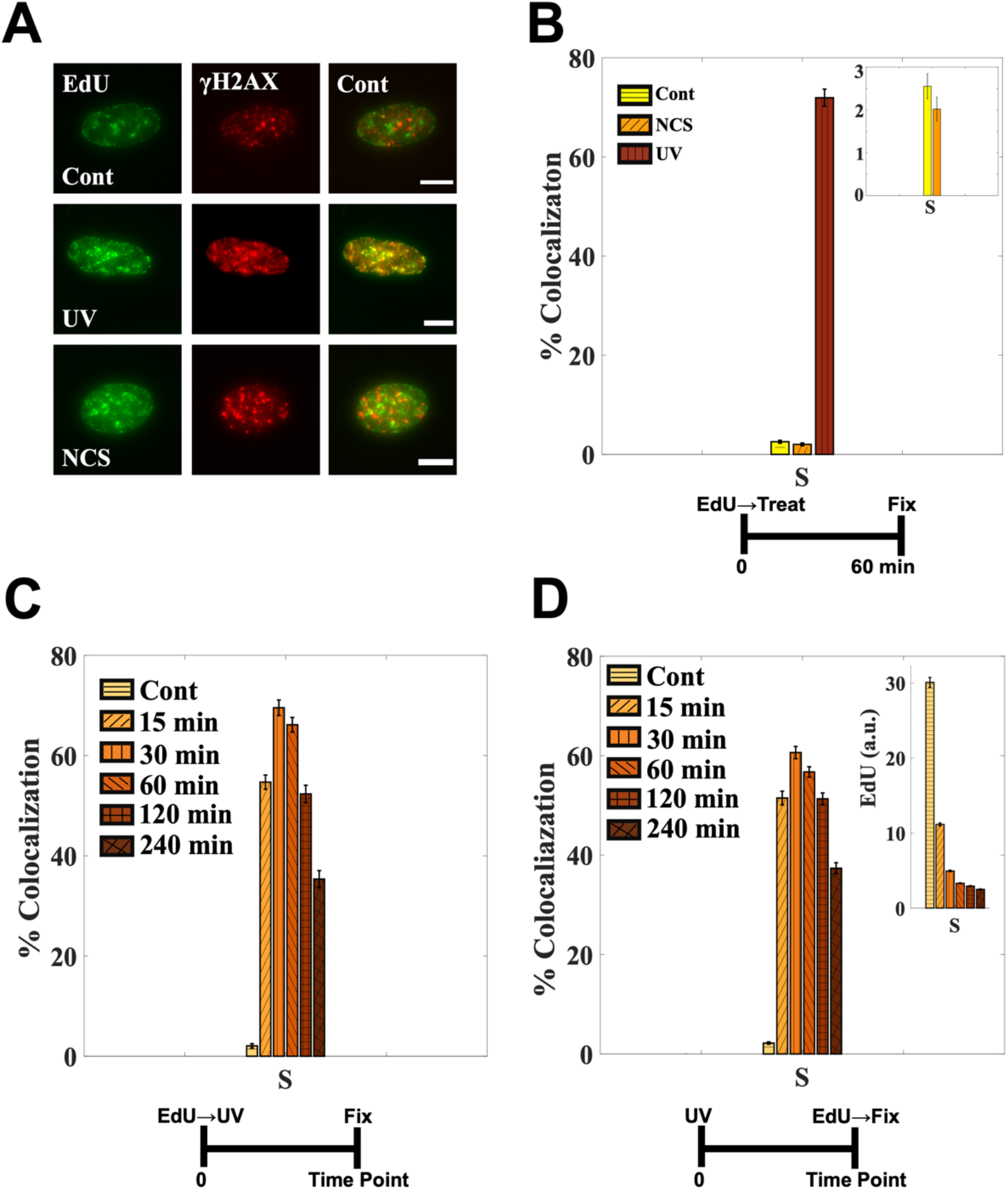
γH2AX colocalizes with the sites of active replication on UV irradiation. (A) Cells were labelled with EdU prior UV or NCS treatment. (B) More than 70% of EdU foci colocalizes with γH2AX foci. The colocalization is less than 3% for control and NCS-treated cells (inset). (C) γH2AX starts to accumulate at sites of active replication within a few minutes of UV exposure as reflected in the percentage of EdU foci colocalizing with γH2AX foci. The colocalization peaks at around 30 minutes after which it starts to go down. (D) Cells were labelled with EdU post-UV, right before fixation. As seen from more-or-less intact colocalization between EdU and γH2AX for the same time points as (C), γH2AX accumulation does not inhibit EdU incorporation altogether but slows it down (inset). See also Supplementary Figure S1. Cells were irradiated with 10 J/m^2^. Scale bar: 10 μm.

To investigate how this localization of γH2AX at the replication forks affects the replication itself, we labelled the cells with EdU post UV but immediately before fixation—unlike in the previous cases where the cells were labelled with EdU right before UV to mark the S phase cells and replication forks at the time of UV irradiation. We observed that EdU was still incorporated at the sites of γH2AX albeit at slower rates as inferred from the intact colocalization between EdU and γH2AX and the lower levels of EdU in those cells (Figure 4D and Supplementary Figure S1). This slowing down of replication could be because of lesion bypass synthesis of DNA on leading and lagging strands. Lesion bypass DNA synthesis are marked by large stretches of single-stranded DNA (ssDNA) which initiates DNA damage repair signalling and recruits DDR proteins [32, 33]. In fact, we observed that most of the S phase cells at the time of UV irradiation did complete the replication and were arrested in the G2 phase of the cell cycle as discussed next.

### S phase cells at the time of UV do not develop proportionately as many DSBs at later time points as indicated by their γH2AX levels

To investigate how the elevated levels of γH2AX affect the S phase cells at later time points, we labelled the cells with EdU prior to UV and let them recover for 24 to 48 hours. We observed that most of the S phase cells at the time of UV irradiation completed the replication of their DNA and stayed arrested in G2 phase even after 48 hours as reflected by their respective EdU staining patterns and cell cycle distributions (Figures 5A, 5B, 5C, 5D and Supplementary Figure S2). Also, the S phase cells at the time of UV irradiation did not develop as many DSBs at later time points as expected from the elevated γH2AX levels observed a few hours post UV (Figures 5C and 5D). Though the cells in G1 phase at the time of UV irradiation showed very few DSBs at later time points (Figures 5C and 5D), those cells which had exited the G1 arrest by then and had started replicating their DNA developed as many DSBs as the ones which had been in the S phase at the time of UV irradiation (Figures 5E, 5F and Supplementary Figure S3). In fact, all the three phases seemed to have similar numbers of pre-apoptotic cells as apparent from the number of cells showing homogeneous pan-nuclear γH2AX staining (Figures 5C, 5D, 5E and 5F). Homogeneous, pan-nuclear γH2AX staining is considered as a marker of pre-apoptotic cells as reported in a previous study [14].

**Figure 5:**
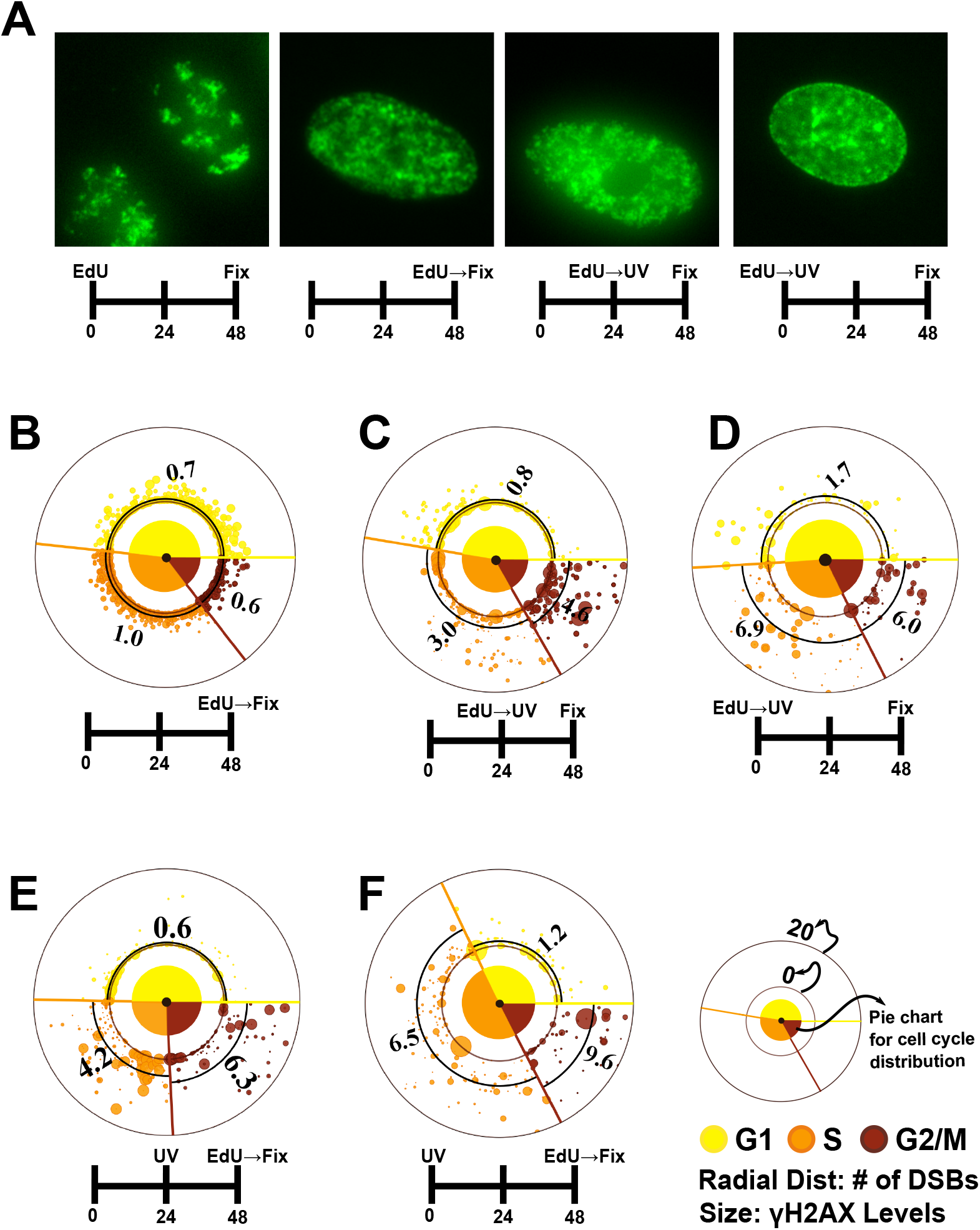
S phase cells at the time of UV irradiation do not have proportionately as many DSBs as indicated by their γH2AX content. (A) EdU staining pattern for control cells post 48 hours is strikingly different from that of UV treated cells which is very similar to that observed for cells freshly labelled with EdU showing that UV-treated cells have not gone through a mitosis and are arrested (see also Supplementary Figure S2). (B, C, D, E, F) Torus plots comparing of number of DSBs and cell cycle distributions for different cases (please see the legends in the figure). DSBs are quantified as described in Figure 3. The numbers on the plots are average DSBs in the respective cell cycle phases as also marked by thick black arcs. The large spots close to the origin represent apoptotic cells. (C, D) The population labelled with EdU right before UV irradiation have their cell cycle distributions very much similar to that of control cells (B) post 24 and 48 hours. It again shows that most of the cells have been arrested and have not gone through a mitosis even at 48 hours post UV. Also the S phase cells in these populations (which were the S phase cells at the time of UV irradiation) do not have as many DSBs at the indicated time points as suggested by the levels of γH2AX. (E, F) The cells here are labelled with EdU post UV, right before fixation. The S phase cells here are those cells which have exited the G1 arrest. These S phase cells, on the average, show similar number of DSBs as the cells which were in the S phase at the time of UV (C and D) showing that UV causes similar levels of damage to all the cells in a population irrespective of their cell cycle phases (see Supplementary Figure S3). Cells were irradiated with 10 J/m^2^.

Taken together the above observations suggest that the UV irradiation does not selectively cause higher levels of damage in the S phase cells; rather it has equitable effects on all the cells in all the phases of the cell cycle.

## Discussion

Ultraviolet radiation is a common environmental mutagen which makes it very important to study UV-induced DNA damage and the responses that it elicits. UV is known to induce replication stress in dividing cells by actuating the dimerization of adjacent pyrimidines that leads to distortions in DNA structures which, if unrepaired, can eventually lead to DSBs— most lethal of all DNA lesions [34]. Most kinds of DNA lesions lead to the phosphorylation of a histone variant H2AX at serine 139 by the master transducers of DNA signals ATM and ATR—a reason why γH2AX is considered a definitive marker of DNA damage. UV exposure, in particular, leads to a sharp peak of γH2AX in the S phase cells which are actively replicating their DNA. This has led to the idea that S phase cells in a population are most vulnerable to genotoxic insults. In this report we systematically study the hypothesis by investigating the nature of γH2AX in the S phase cells after DNA damage.

We first show that not all genotoxic treatments lead to a γH2AX peak in the S phase of the cell cycle: UV treatment causes γH2AX to peak in the S phase but NCS, a DSB-causing agent, does not. We also show that UV-induced S phase peak in γH2AX levels is not reflected in the levels of primary (in terms of CPDs) or secondary DNA damage (in terms of DSBs) caused by UV. This shows that it is the damage response and not DNA damage itself, which peaks in the S phase post UV irradiation. We also find that the increase in γH2AX after UV irradiation in the S phase within a population does not square with the decrease in global transcription—a known measure for the extent of DNA damage, especially of that induced by UV exposure, inside a cell [27]. We also observe that γH2AX colocalizes with the sites of active replication and that those sites continue incorporating EdU even 4 hours past UV irradiation, although at slower speeds. This hints at underlying mechanisms with which cells deal with stalled replication forks from bulky DNA lesions via either translesion synthesis or re-priming events downstream of those lesions in both leading and lagging strands of DNA [32, 33]. Such events are reported to produce large stretches of ssDNA at replication forks which initiate DNA damage signalling by recruiting DDR proteins [32, 33]. These observations suggest that the colocalization of γH2AX foci in the S phase cells post UV exposure mark damage response rather than the extent total damage in those cells. The inference is further strengthened by the observation that those cells in the S phase at the time of UV irradiation do not develop proportionately as many DSBs at later time points as expected from their γH2AX levels. In fact, the number of DSBs induced by UV at later time points seems to depend on the DNA content: it increases with the increase in DNA content as moving from G1 to S to G2 phases of the cell cycle similar to what is observed in cells treated with NCS. Taken together, our results suggest two things: 1) vulnerability to the exogenous sources of DNA damage depends more on the DNA content than on the cell cycle stages and 2) the γH2AX foci in the S phase after UV irradiation are not the sites of DNA damage—they colocalize with the active replication forks at the time of UV irradiation. This also suggests that the sites of active replication post UV irradiation do not correspond to DNA lesions in congruence with a recent study on the yeast *S. cerevisiae* system showing that UV-induced DNA lesions are spatially and temporally separated from the active replication forks [35].

Although these results have been shown just in the context of UV-induced replication stress, they might also hold true for replication stress induced by other genotoxic agents such as camptothecin or 4NQO or cisplatin where similar observations regarding γH2AX have been reported [8–10, 36]. This poses new questions as to the roles of γH2AX in the maintenance of genome stability which hitherto has been confined just to the mediation of DNA repair inside a cell. It also highlights the need to look at chromatin modifications such as γH2AX with the broader view involving the gamut of processes the chromatin modifications control. In future it will be interesting to study how γH2AX is involved at maintaining stability of replication sites after UV irradiation, although it has been questioned if the presence of H2AX is necessary for cell survival [37]. It would shed light on the possible roles and importance of γH2AX in fork stability in the mammalian system analogous to that recently seen in the yeast *S. pombe* [38].

## Materials and Methods

### Cell culture and treatments

A549 cells were used for all the experiments. Cells were grown in DMEM/F12 (Gibco) supplemented with 10% FBS (Gibco) without antibiotic. Cells were tested negative of any bacterial contamination including mycoplasma. Cells were passaged every ~4 days and were let to grow for at least 24 hours before starting any experiments.

Cells were labelled always with 10 μM 5-Ethynyl-2’-Uridine (EdU) for 15 minutes before or after UV as the case may be, to mark true S phase cells.

For UV treatment Analytik Jena UVP crosslinker CL-1000 was used. The instrument was set to the appropriate dosage in J/m^2^ before irradiation. The cells in plates were washed twice with PBS before irradiating them with 254 nm UV light. The irradiation was done in the absence of any liquid medium in the plates.

Neocarzinostatin was used at 1.6 μg/ml concentration. The cells were treated with the chemical-containing medium for 2 minutes after which the cells were washed twice with PBS and fresh medium was added to the plates.

Cells were let to recover after all treatments for 60 minutes unless mentioned otherwise. For all the experiments, including long time (24 and 48 hours) point experiments, the cells were plated at the same time and treated exactly the same but for their respective experimental conditions.

### Immunofluorescence and click chemistry

Cells were fixed with 4% PFA in PBS for 15 minutes followed by permeabilization with 0.3% Triton-X 100 in PBS for 10 minutes. For immunofluorescence combined with click chemistry, permeabilization was followed by click chemistry detection of EU or EdU followed by the usual immunofluorescence as described below.

For immunofluorescence-based detection of protein levels, cells were blocked for nonspecific binding by using Block – a 5% solution of bovine serum albumin (BSA, Sigma A2153) in PBS – for 30 minutes at room temperature (RT) followed by labelling with the primary antibodies in Block for 60 minutes at RT. The cells were then washed twice with PBS and were labelled with the secondary antibodies in Block for 60 minutes at RT.

The protocol for click chemistry was adopted from [39]. In brief, cells were equilibrated with TBS buffer before the addition of click reaction cocktail (CRC) to label the cells with azide dyes. 1 ml of CRC was prepared thus: 730 μl of 100 mM Tris (pH ~ 8.5) + [15 μl of 100 mM CuSO_4_ + 41 μl of 50 mM THPTA (tris-hydroxypropyltriazolylmethylamine (Sigma 762342)) ligand + 1 μl of 6 mM Azide dye] + [100 μl of 10% glucose in 10% glycerol + 4 μl of 20K U/ml stock of catalase (Sigma C3515)] + 8 μl of 500 U/ml glucose oxidase (Sigma G2133)] + 100 μl of 1 M sodium ascorbate.

The order of reagent addition was important. Reagents in the square brackets were prepared separately in two 1.5 ml tubes. The tube containing THPTA, CuSO_4_ and Azide dye was let to sit for at least 5 minutes before Tris was added to the tube. After which the content of the other tube was added to the tube containing THPTA. Finally sodium ascorbate was added.

Once CRC was prepared TBS in plates was replaced with CRC. The labelling was performed for 15 minutes after which the cells were washed with PBS and then with 100 mM EDTA in PBS for 5 minutes to remove excess Cu ions from the cells. After which the cells were treated for the usual immunofluorescence.

Lastly the cells were labelled with 1 μg/ml Hoechst 33342 in PBS for 10 minutes.

For measuring global transcription, cells were labelled with 5-Ethynyl-Uridine (EU) immediately after UV irradiation for 30 minutes. EU gets incorporated into actively transcribing RNAs and marks the translational activity inside the cells which can later be detected using click chemistry. EU (CLK-N002-10) and EdU (CLK-N001-100) were procured from Jena Bioscience. The stock solutions were made in PBS.

For CPD staining, cells were denatured right after permeabilization with 2N HCl at 37°C for 15 minutes. Cells were then washed once with PBS and then neutralization was performed with 0.2 M Borax buffer with pH ~ 8.5 for five minutes at RT. After which Click chemistry and IF followed as described above. DNA denaturation required longer Hoechst staining achieved by adding Hoechst directly to the secondary antibodies solution.

### Antibodies used

Anti-γH2AX from Merck (05-636) at 1:500 dilution, Anti-53BP1from Abcam (ab175933) at 1:1000 dilution and Anti-pATM from Abcam (ab36810) at 1:250 dilution. CPD antibodies from Cosmo Bio (TDM-2 clone, CAC-NM-DND-001) at 1:250 dilution. Highly cross-adsorbed Alexa Fluor-labelled secondary antibodies raised in goat from Invitrogen were always use at 1:500 dilution.

### Fluorophores

The fluorophores were kept constant across the experiments: Click chemistry was always performed with azide Alexa Fluor 488. Mouse antibodies (which include Anti-γH2AX) were always detected with Alexa Fluor 546 and rabbit antibodies (which include Anti-53BP1) with Alexa Fluor 594. DNA was stained with a minor groove-binder Hoechst 33342.

### Microscopy

Imaging was always performed with freshly-prepared 0.1% solution of p-phenylene diamine (PPD) in PBS as a mounting medium to reduce photobleaching of Hoechst 33342.

Images were acquired at 14 bit resolution using the fully automated Olympus IX83 microscope on a Retiga 6000 (QImaging) camera. All images were taken using a 40X, 0.75 NA air objective. Four planes, 2 μm apart, were taken for every field. Filter sets from Olympus and Chroma Technology Corp (49309 ET – Orange#2 FISH and 49310 ET – Red#2 FISH) were used to prevent any bleed through among the fluorophores used.

### Data and image analyses

Cells were first staged in the cell cycle based on their DNA content using an automated Matlab routine developed earlier [8]. Once staged, the cells were later redistribute cross the cell cycle stages based on their EdU intensities to identify true S phase cells. For experiments where it was not possible to combine EdU-based staging of cells, cells were staged based just on their DNA contents (Figure 3). All the analyses included at least 500 cells in total for all the three cell cycle phases.

All image and data were analysed using Matlab. For graphs Matlab and Python 3 were used. All codes and programs used in the study are available on request.

## Author contributions

AM, SD, SR conceived and designed the experiments. SD performed and analysed the experiments. SD, SR, AM interpreted the data. SD, AM wrote the manuscript. AM monitored the project and associated grants.

## Conflict of interest

The authors declare that they have no conflict of interest.

## Funding

This project was funded by intramural funds at TIFR Hyderabad from the Department of Atomic Energy (DAE), and partially by a DST SERB Early Career Research Award (ECR/2016/000907) to AM. SR was supported by DST SERB National Post-Doctoral Fellowship (N-PDF) scheme (PDF/2016/002781).

## Acknowledgements

AM and SD thank Dr. Tamal Das for access to the UV crosslinker.

---

**Supplementary Figure S1:**
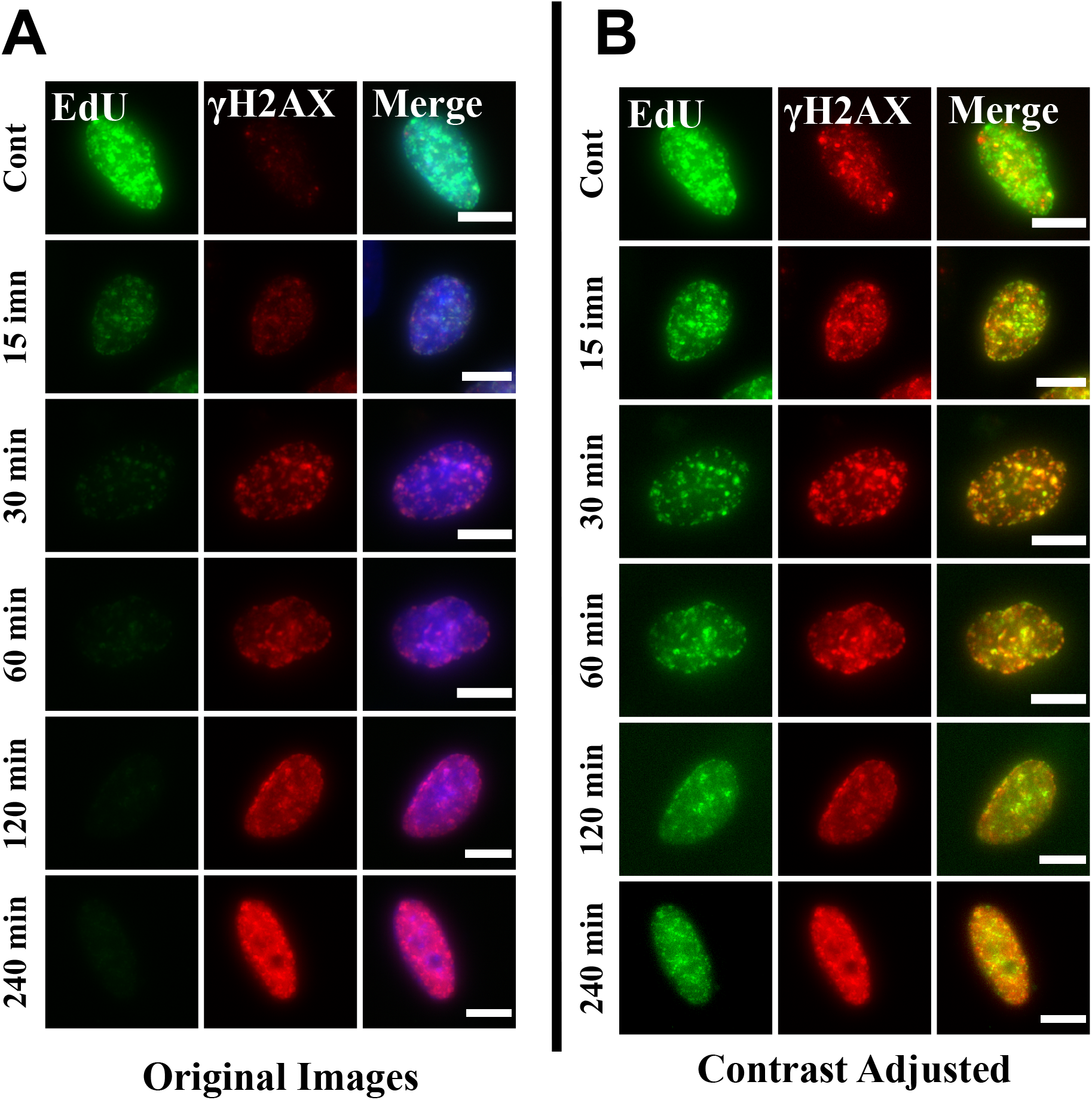
γH2AX accumulation at the sites of active replication after UV does not stop replication (Related to Figure 4) Cells were labeled with EdU just before fixation. (A) EdU incorporation goes down with time after UV exposure. (B) Colocalization between γH2AX and EdU is intact as can be seen for the same cells as in (A), but contrast adjusted to show weak signals. This colocalization implies that UV exposure slows down the replication in cells but does not stop it altogether. Cells were irradiated with 10 J/m^2^. Scale bar: 10 μm.

**Supplementary Figure S2:**
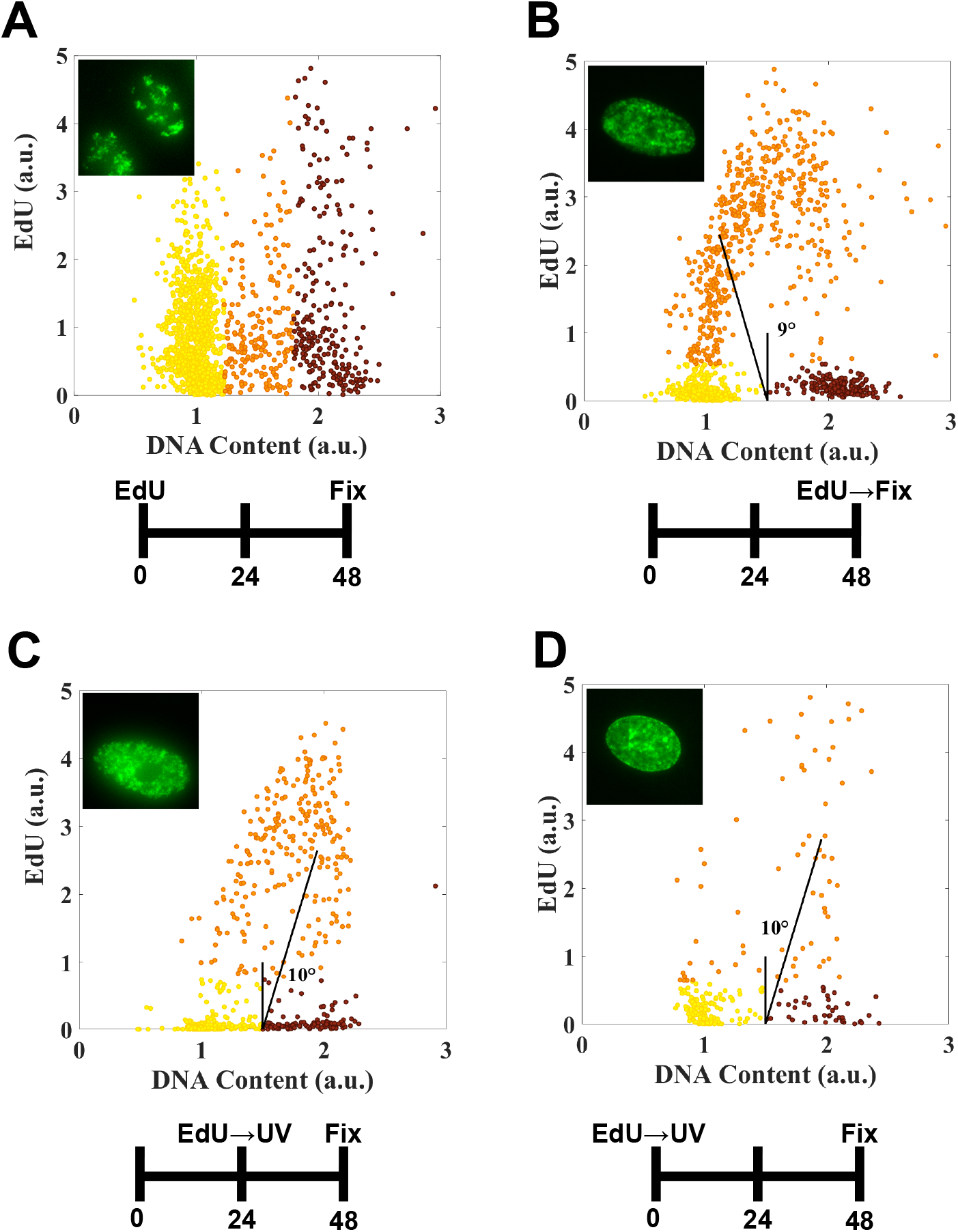
S phase cells at the time of UV irradiation completes replication and arrest in the G2 phase of the cell cycle (Related to Figure 5) (A) Cells labelled with EdU 48 hours prior to fixation. Cell cycle distribution of EdU in this population does not show an arch as seen in that for freshly-labelled EdU cells as shown in (B). Inset shows the staining pattern for these cells forming distinct patches. (B) Cell cycle distribution for EdU in freshly-labelled EdU cells. The DNA content mode of the S phase cells is closer to that of G1 phase cells as indicated by the line making a 9° angle towards left from the vertical. (C, D) Cells were labelled with EdU right before UV irradiation and let to recover for 24-48 hours. The EdU distributions and staining patterns are strikingly similar to that for freshly-labelled EdU cells. The DNA content modes of S phase cells in these population have shifted closer to that of G2 phase cells as indicated by the lines forming 10° angles towards right with the vertical in both the cases. This shows that most of the S phase cells have completed the replication and are arrested in G2 phase of the cell cycle even at 48 hours post UV. The cells for all these experiments were plated at the same time and were treated exactly the same but for their respective experimental conditions; this explains the fewer cells observed in UV-treated populations. Cells were irradiated with 10 J/m^2^.

**Supplementary Figure S3:**
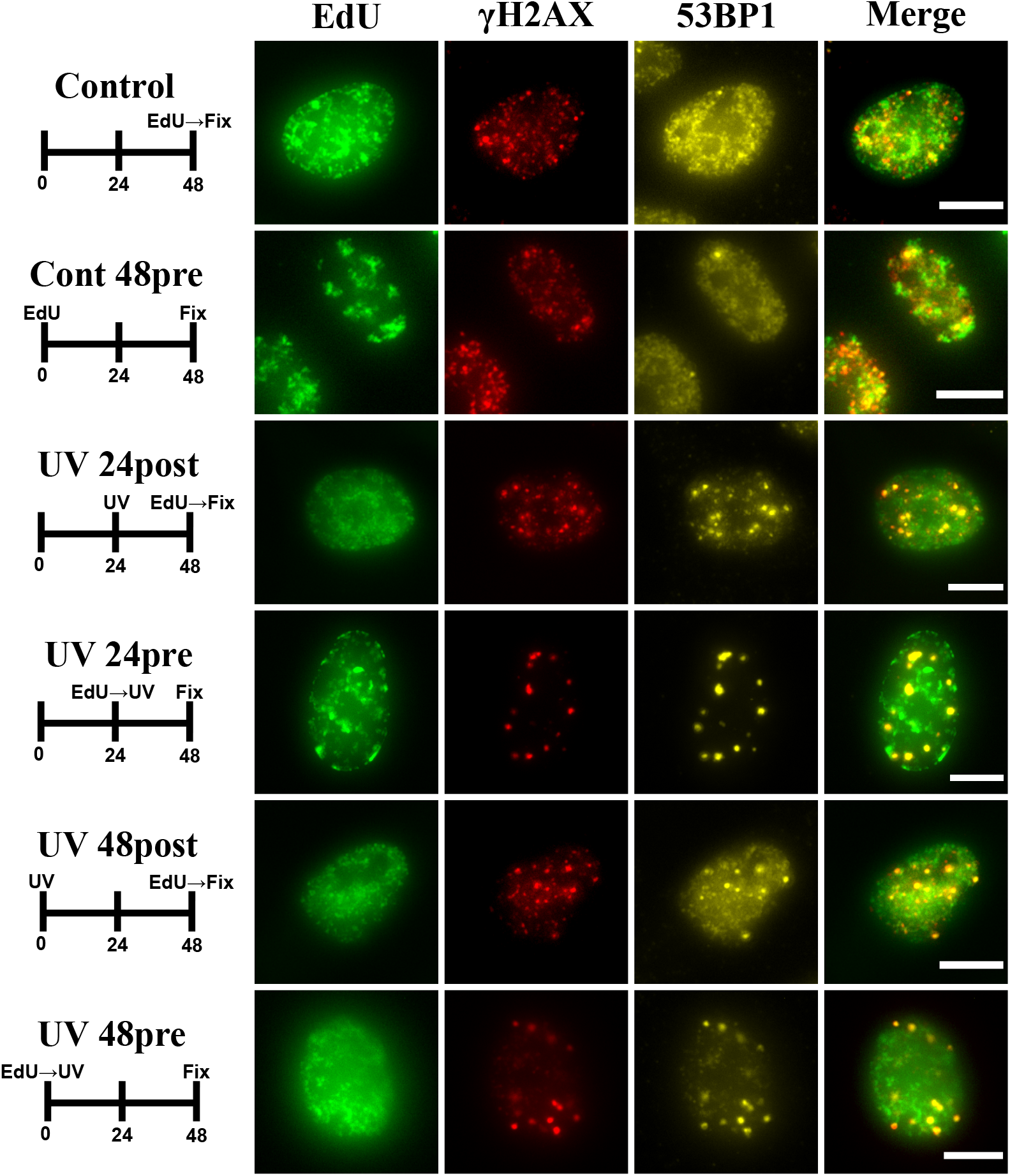
S phase cells at the time of UV irradiation do not have proportionately as many DSBs as indicated by their γH2AX content (Related to Figure 5) S phase cells are shown for all the different cases. Colocalization of γH2AX and 53BP1 marks DSBs. S phase cells at the time of UV have as many DSBs at later timepoints (4^th^ and 6^th^ rows) as the cells which were in G1 phase but now have exited G1 arrest and started replicating (3^rd^ and 5^th^ rows). The quantification for this observation is shown in Figure 5. The images are contrast-adjusted to aid visualization. Cells were irradiated with 10 J/m^2^. Scale bar: 10 μm.

## Notes

### Competing Interest Statement

The authors have declared no competing interest.

### Summary of Updates

Addition of a new experiment (Figure 2).

